# On mechanisms of light-induced deformations in photoreceptors

**DOI:** 10.1101/2020.01.08.897728

**Authors:** Kevin C. Boyle, Zhijie Charles Chen, Tong Ling, Vimal Prabhu Pandiyan, James Kuchenbecker, Ramkumar Sabesan, Daniel Palanker

## Abstract

Photoreceptors in the retina convert light into electrical signals through a phototransduction cycle that consists of multiple electrical and biochemical events. Phase-resolved optical coherence tomography (pOCT) measurements of the optical path length (OPL) change in the cone photoreceptor outer segments after a light stimulus (optoretinogram) reveal a fast, ms-scale contraction by tens of nm, followed by a slow (hundreds of ms) elongation reaching hundreds of nm. Ultrafast measurements with a line-scan pOCT system show that the contractile response amplitude increases logarithmically with the number of incident photons, and its peak shifts earlier at higher stimulus intensities.

We present a model that accounts for these features of the contractile response. Conformational changes in opsins after photoisomerization result in the fractional shift of charge across the disk membrane, leading to a transmembrane voltage change, known as the early receptor potential (ERP). Lateral repulsion of the ions on both sides of the membrane affects its surface tension and leads to its lateral expansion. Since the volume of the disks does not change much on a ms time scale, their lateral expansion leads to an axial contraction of the outer segment. With increasing stimulus intensity and resulting tension, the area expansion coefficient of the disk membrane also increases as thermally-induced fluctuations are pulled flat, resisting further expansion. This results in a logarithmic saturation of the deformation and a peak shift to earlier with brighter stimuli. Slow expansion of the photoreceptors is explained by the influx of water due to osmotic changes during phototransduction. Both effects closely match measurements in healthy human volunteers.

## Introduction

Retinal photoreceptors convert light into electrical signals through a phototransduction cycle that operates across multiple time scales, from milliseconds to seconds. Understanding the resulting electrical and biochemical activity throughout this response is very important for comprehending visual function in general, and diagnosis of its pathology in particular. Until now, electrical signals in photoreceptors and other retinal cells have been measured using intracellular or extracellular electrodes [1]–[3]. Recently, phase-resolved imaging of photoreceptors demonstrated significant changes in the optical path length (OPL) in response to flash stimuli [4]–[6]. Similarly, nm-scale OPL changes accompanying the action potential have been observed in spiking neurons ex-vivo, and have been shown to faithfully reproduce changes in the cell potential [7]–[9]. Interferometric imaging of such nm-scale changes over ms time scales offers a non-invasive and label-free alternative to current electrophysiological methods, which may allow in-vivo studies of neurobiology, especially retinal physiology.

To relate the observed mechanical deformations of cells to the underlying physiological processes, the coupling mechanisms must be known. Recently, we demonstrated that nm-scale deformations in neurons and other electrogenic cells can be explained by the voltage dependence of the membrane tension [8], [10]. Changes in the membrane tension necessarily cause a deformation to rebalance forces across the structure, whether it’s a whole cell or an organelle. The charged membranes in photoreceptors are no exception and should similarly link electrical activity to deformations.

Here we study the mechanisms underlying the changes in photoreceptor outer segments following pulsed stimuli, as observed in human eyes using phase-resolved optical coherence tomography (pOCT). We present a model of the light-induced deformation that fits the measured data across multiple stimulus intensities and discuss how such a model can aid in characterizing the phototransduction cycle or the mechanical properties of the disk membranes. We also discuss how the model can help enhance the sensitivity of future recordings.

## Observed OPL changes in human retina

Phase-resolved optical coherence tomography (pOCT) recordings in the living human eye demonstrated significant optical path length (OPL) changes in the outer segments of photoreceptors in response to light stimuli. It begins with a fast, ms-scale contraction by a few tens of nm, followed by a much slower (hundreds of ms) and much larger (hundreds of nm) elongation [11], [12]. These deformations are assumed to be linked to the underlying electrical activity in the photoreceptors and have accordingly been termed the optoretinogram, analogous to the electroretinogram. This phenomenon has been used to classify cone types in-vivo [4]. The slow expansion has been attributed to water influx due to the osmolytes produced in the phototransduction cascade [5], [12], but the mechanism of the fast contractile response remains unknown.

In the current study, we used a fast line-scan pOCT system, which enabled capturing the dynamics of the fast response, and allowed exploration of the underlying mechanism. Compared to the point-scan and full-field OCT systems used in the past [5], [13], a line-scan system captures an entire b-scan in each frame of the camera, and thereby achieves sub-ms time resolution, while recording structural information sufficient for the image registration. This is achieved by recording the spatial and spectral components of a b-scan across the two axes of a 2D imager [11]. The line-scan pOCT system applied here uses a superluminescent diode (*λ*_0_ = 840 *nm*, Δ*λ* = 50 *nm*) for the OCT path and a 528 ± 20 *nm* LED for retinal stimulation. To enable single-cone resolution, it includes adaptive optics implemented with a 97-element deformable mirror with pupil diameter of 13.5 mm and a custom Shack-Hartmann wavefront sensor at 980 ± 10 *nm*. For high resolution measurements of the early response, it achieves an imaging field of 2 × 0.6 *deg* on the retina, 5 − 7 *deg* temporal to the fovea at a volume rate of 324 Hz. The early response recordings were captured by averaging 500 b-scans per volume at a rate of 16.2 kHz, which can be reduced to an average of two b-scans for a maximum temporal resolution of 123 μs. Such speeds are sufficient for revealing the details of the fast response in the cone outer segments, which is a 5-40 nm contraction over less than 5 ms, preceding the slow swelling response. System sensitivity was 92 dB with phase sensitivity of 4 mrad at 50 dB SNR, where 1 *mrad* ≈ 0.07 *nm* at *λ*_0_ = 840 *nm*.

## Early receptor potential

We assume that the observed ms-fast decrease of the OPL through the photoreceptor outer segments is related to the early receptor potential (ERP)—a ms-fast electrical signal following a bright stimulus, which can be recorded in cone photoceptors via patch clamp [1], [2]. The ERP is attributed to the charge transfer across the cell membrane associated with the conformational change of the opsin (photoisomerization) embedded in that membrane. This ERP signal is distinct from the late receptor potential (LRP), which corresponds to changes in the cell potential associated with closing of the ion channels in outer segments following the phototransduction cascade [1], [2]. As illustrated in Figure 1, the change from 11 cis- to all-trans conformation of the opsin during isomerization effectively results in the movement of a single charge by approximately 12% across the cell membrane. This corresponds to a 10^−10^ *V* change in the membrane potential per photoisomerization. With an increasing number of incident photons, the ERP first increases linearly, but then asymptotically (exponentially) reaches the saturation level as the 10^8^ opsin molecules in the cone photoreceptor are increasingly bleached, reaching a saturated ERP level of *U* = −14.3 *mV* [2].

**Figure 1.**
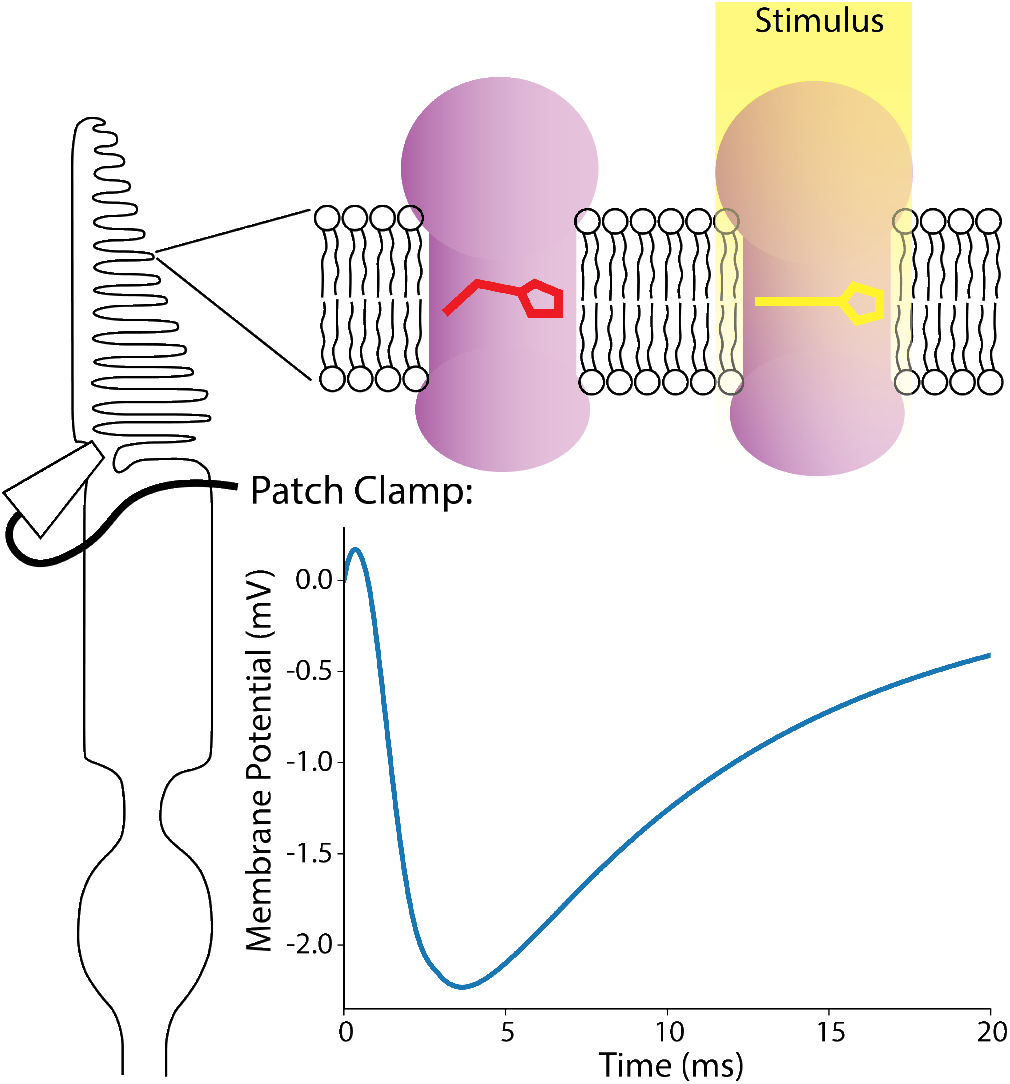
Patch clamp measurements of the cone outer segment demonstrate the transmembrane potential transient following a 1 ms flash, caused by the conformation change of the opsins embedded in the disk membranes. Each photoisomerization induces a 10^−10^ *V* potential change, leading to a characteristic potential change up to a few mV in the outer segment, with its time course shaped by the time constants of the various steps in the phototransduction cycle.

## Voltage-dependent membrane tension

Rapid cellular deformations accompanying the changes in the transmembrane potential have been observed in multiple cell types, including the recent full-field interferometric imaging of HEK cells and neurons in culture [7]–[10], [14]. These measurements have established a linear relationship in typical cells between the membrane displacement and changes in cell potential, linked by the dependence of the membrane tension on transmembrane voltage.

The shape of biological cells is determined by the balance of intracellular hydrostatic pressure, membrane tension, and strain exerted by the cytoskeleton [15]. The membrane tension includes a contractile component from the lipid bilayer and the lateral repulsion of ions in the Debye layer on both the intracellular and extracellular sides of the membrane:

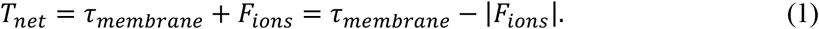

The transmembrane potential changes due to movement of ions across the cell membrane, which affects the charge density in the layers of mobile ions along the inner and outer surfaces of the membrane. Change in the lateral repulsion of the ions, in turn, affects the membrane tension, and the cell must rebalance the forces via a deformation. The lateral repulsion of ions can be expressed as a function of the charge density *σ* and ionic strength *n* on the inside, *in*, and outside, *ex*, of the membrane, as well as the trans-membrane voltage *V* [7]:

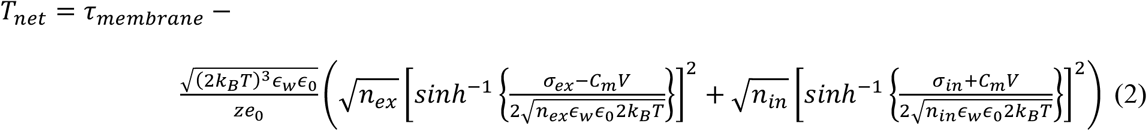

Here, k_B_ is Boltzman’s constant and T is the absolute temperature, ϵ_w_ and ϵ_0_ are the relative permeability of water and the permittivity of free space, z is the number of valence ions in the solution, e_0_ is the electronic charge, and C_m_ is the capacitance of the membrane between the Debye layers. The membrane tension increases by about 0.1 *mN*/*m*/*V*, or 10 μN ∙ m^−1^ for a 100 *mV* depolarization, which is a decrease of trans-membrane voltage [8]. Increased surface tension of the cell membrane leads to deformation of a cell toward minimization of its surface area, i.e. becoming more spherical, which would be the case during the action potential [10].

## Quasi-static model

During hyperpolarization of the photoreceptor under illumination, the lateral repulsion of ions *F_ions_* in the charge layers increases, and hence the disk membrane area will expand. If the disk volume during a few ms after the flash remains constant, widening of the disks will flatten them, leading to shortening of the outer segments, as illustrated in Figure 2.

**Figure 2.**
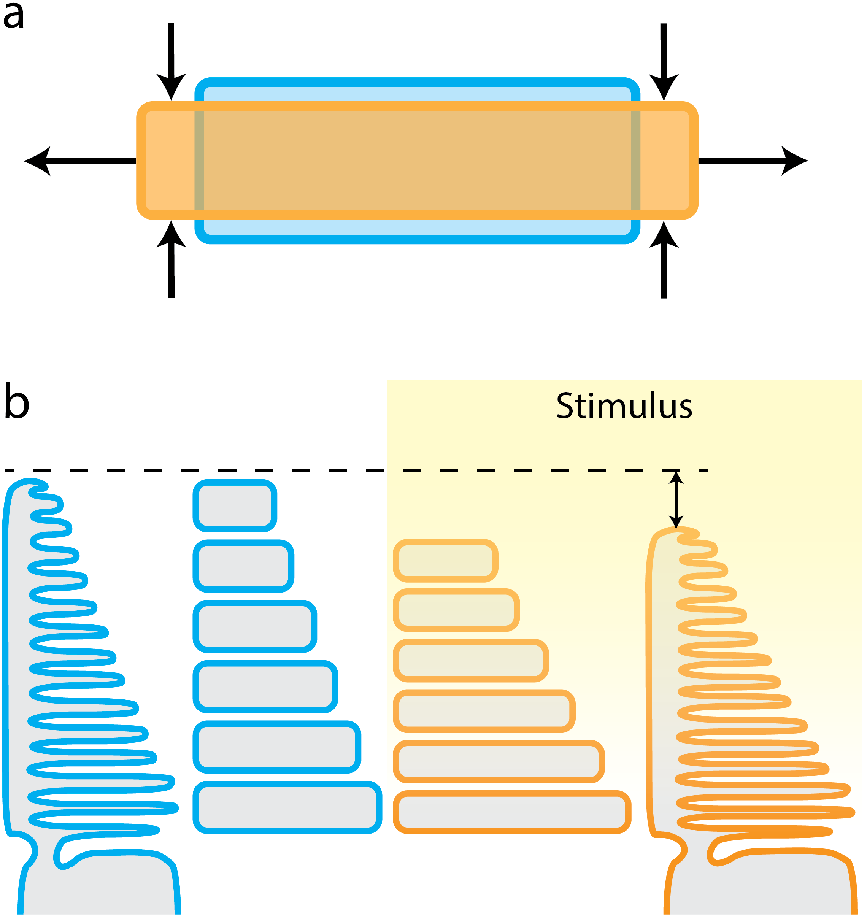
The deformation model of the photoreceptor outer segments assumes (a) the membrane expansion and subsequent flattening of the fixed-volume disks following the stimulus, which result in (b) the shrinkage of the outer segment that contains a stack of approximately one thousand disks.

The disk membranes in photoreceptor outer segments have very low tension since they are formed as part of a blebbing process and contain no actin cortex [16]–[18]. Due to this lack of structure, they lay flat in stacks that are held together by inter-disk proteins [19]. The low tension allows the membrane to undulate due to thermal fluctuations [20], and even small forces can easily pull the undulations out, leading to lateral expansion of the membrane, as shown in Figure 3a. The membrane area expansion coefficient at low tension is defined by the bending modulus *k*_*c*_ = 0.5 × 10^−19^ *N* ⋅ *m* [20], [21], while at high tension, when the undulations are flattened-out, the area expansion modulus of a flat membrane *K*_*A*_ = 0.2 *N*/*m* dominates [22]. The apparent modulus in-between these two extremes varies as a function of tension 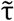 [23]:

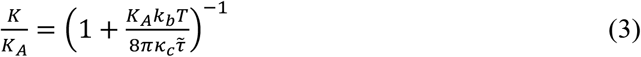

At low tensions, *K* scales approximately linearly with 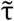:

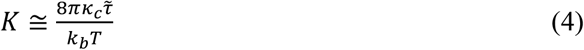

**Figure 3.**
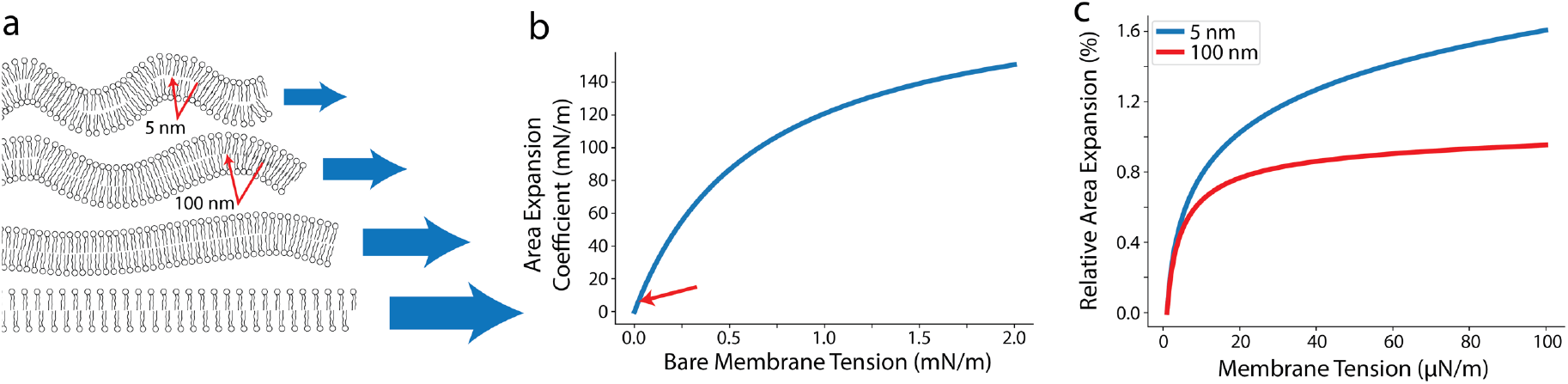
Saturation of the deformations due to the increasing expansion modulus of the disk membranes. (a) At low tensions, the disk membranes contain thermally induced fluctuations that require little force to pull out, depending on the smallest bending radius as labeled, but at higher forces the intermolecular bonds of the membrane begin to dominate. (b) The resulting area expansion coefficient increases with tension, and in the small range of tensions relevant to the outer segment disks, indicated by the red arrow, can be approximated as a linear function of tension. (c) The relative area expansion thus saturates logarithmically over a 100*μN*/*m* tension change. The rate of saturation depends on mechanical parameters of the membrane, notably including the minimum bending radius illustrated here, where 5 nm is the size of an individual lipid head, while the larger minimum radii could be defined by the much larger opsins embedded in the membrane.

The range relevant for the physiological conditions of the disk membrane is very narrow, as indicated by the red arrow in Figure 3b. This linear relationship between the area expansion modulus and tension suggests a logarithmic relationship between the integrated expansion and applied tension in that range of interest. Following this intuition, and based on the thermodynamic equilibrium considerations, Marsh [23] derives the normalized area expansion of a lipid membrane as:

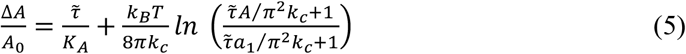

where *A* is the size of the membrane patch (in this case the area of the photoreceptor disk face), *A*_0_ is the area at zero observable tension 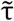, and *a*_1_ is the smallest bending feature size (area of a lipid head in a simple lipid bilayer membrane). In the case of a membrane densely filled with rhodopsin (typically making up 50% of the membrane area), *a*_1_ is likely not limited by the lipid head width, but rather by the characteristic size of the opsin nanodomains embedded in the disk membrane, which are about 25 nm in size [24]. Figure 3c illustrates the saturation behavior of the area expansion with increasing tension for different values of *a*_1_.

Considering a fixed volume of the disk as a cylinder *V* = *A* ∙ *z*, the change in thickness of the disk Δ*z*/*z* to accommodate this area expansion is:

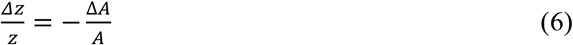

Due to the double pass of light reflected at the bottom of the outer segment, the corresponding change of the OPL is a function of its total height and the index of refraction:

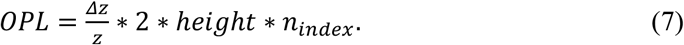

With light intensities well below saturation of absorption (<15% bleaching of the opsins), the membrane potential change in the disk does not exceed a few mV and scales approximately linearly with the number of photons. This corresponds to about 1/100^th^ *nm* OPL decrease per disk, but in an outer segment of 10 *μm* in height, made up of a stack of about one thousand disks, the overall deformation is in tens of *nm* (Figure 2).

As mentioned earlier, the membrane tension increases with the trans-membrane voltage by 0.1 *mN*/*m*/*V*. Driving this tension change with the known time course of the ERP (Figure 1, [2]), we can calculate the area expansion over time for this quasi-static model. To account for the slow swelling attributed to osmotic changes during phototransduction, as well as any tension change from the late receptor potential, we also include a term linearly increasing with time [12], [25]. As shown in Supplementary Figure 1, this quasi-static model provides a poor fit to OPL changes measured in the living human eye. Most notably, this quasi-static model does not allow for the negative peak to shift later in time with less intense stimuli. However, the amplitude of the deformation is close to the observations with reasonable physical parameters, as summarized in Table 1, implying that the general basis of the model is sound.

**Table 1.** Quasi-static model parameters

| Parameter | Fit | Typical |
| --- | --- | --- |
| Slow Response Slope | 1.8e-6 m/s | ~1e-6 m/s |
| Initial Tension | 0.2 $\mu N/m$ | 0.1 - 1 $\mu N/m$ |
| Bending Modulus | 2e-19 N-m | 0.5 - 2e-19 N-m |
| Tension/Voltage (N/m/V) | -0.9 mN/m/V | -0.1 mN/m/V |
| Smallest Patch Size | 10 nm | 25 nm |
| Disk Radius | 3 $\mu m$ | 1-3 $\mu m$ |

## Time-dependent model

To achieve a better fit, we must consider the dynamics of the disk membrane. Since the membrane stiffness K increases with tension, we expect it to behave as a non-linear spring (Supplementary Figure 2). This could account for both the saturation of the peak amplitude, and the faster time to peak with stronger stimuli. Assuming a linear scaling of the area expansion coefficient with tension in the range of interest (Figure 3b), the membrane can be modeled as a spring with a stiffness linearly increasing with the applied force and an offset:

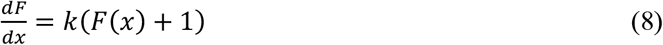

where k is a spring constant we will fit to the data. The restoring force is then an exponential function of the displacement: *F*(*x*) = *e*^*kx*^ − 1, and deformation scales logarithmically with the applied force: 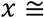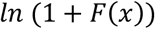, which is analogous to the scaling of the area expansion of the lipid membrane with tension [23]:

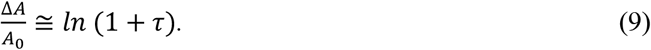

Adding a velocity-dependent viscosity term, the membrane dynamics can be described as following:

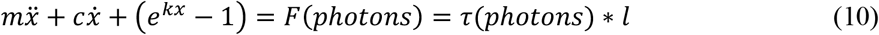

Fitting the parameters (*m*, *c*, *k*, and *l*) to the data yields a very good match of both the amplitude and the time course of the deformation at different stimulus intensities. Figure 4a shows the contributions from the fast response (ERP) and the linear slow response originating in the late receptor potential and osmotic influx of water. The time course of the OPL changes recorded at two stimulus intensities were used to fit the parameters in Figure 4b, with the peak timing now properly accounted for. The same fit parameters were then used to calculate the peak displacement across multiple stimulus intensities, and the resulting dependence matches well to the experimental observations, as shown in Figure 4c.

**Figure 4.**
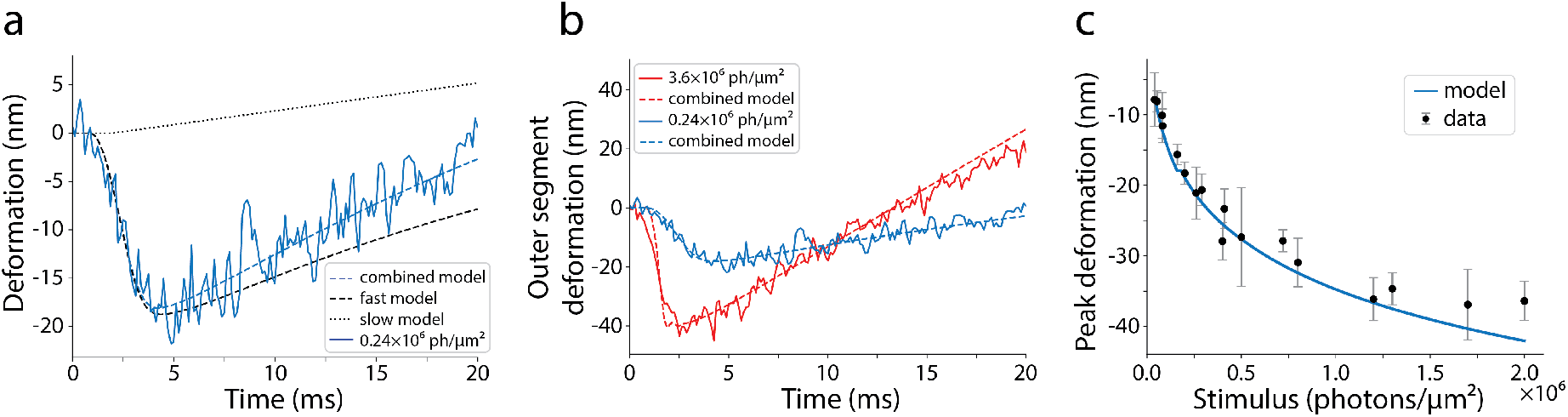
The time-dependent deformation model fits the observations well. (a) The combined model is comprised of two components: the fast contraction caused by the ERP and the linearly rising slow expansion that represents both the LRP and the osmotic swelling of the outer segment. (b) The model parameters are fit to observations at two stimulus intensities, and then the same parameters are used in (c) to predict the peak deformation as a function of stimulus intensity from additional measurements.

## Discussion

The slow response modeled here as a linear function of time (Figure 4) accounts for two effects: the voltage-dependent tension change resulting in membrane expansion due to the late receptor potential (LRP), which begins about 4 ms after the stimulus [2], and the much larger and longer-lasting onset of the osmotic swelling [12]. The phototransduction cascade and the timing of its various stages is well studied [26], so the magnitude and dynamics of the swelling can be compared to these processes. Zhang et al. [12] suggest that the g-protein complex disassociating from the disk membrane after photoisomerization releases transducin into the cytoplasm that could account for part of the osmotic misbalance and the associated influx of water into the outer segment. Comparison of the expected swelling from transducin to observations indicates that other osmolytes seem to contribute to this process as well [11].

A strong ERP signal has only been observed in patch clamp measurements of cones, whose disks are continuous with the outer segment membrane, and hence the transmembrane potential is accessible to the probe patching the cilium [1], [2]. In rod photoreceptors, however, most of the disk membranes are pinched off and stacked within the outer segment membrane. Since the ERP is associated with the translocation of rhodopsin across the disk membranes, very little potential change occurs across the outer segment membrane that the patch clamp has access to in rods. The small signal visible in such measurements is attributed to the minority of rhodopsin embedded in the outer segment membrane as well as the few emerging disks at the base of the outer segment that have yet to pinch off [16]–[18]. However, optical measurements of the fast response, in which mechanical deformations in each independent disk add up, allow measurement of the ERP in rods without the electrical limitations of the patch clamp.

While the quasi-static model presented here estimates the magnitude of the deformations with physiologically correct parameters, and the dynamic model fits the measurements well, we leave it to future work to connect the non-linear spring model with the underlying physical parameters of the membrane in thermal equilibrium. Such a model should account for distributed mass and other physical parameters of the disk membranes, including their base tension, bending modulus and minimum patch size, linked to the size and density of the embedded opsin nanodomains [24]. Such an analytical model would link many mechanical and electrical characteristics of the disk membranes with the observable outer segment deformations.

Characterization of the nm-scale deformations in other electrically active cells, such as HEK cells and cortical neurons, has benefited from a thorough understanding of the underlying mechanisms relating the membrane potential change to mechanical deformation [8], [10]. Cellular responses averaged over many trials provide a template for match-filtering temporal signals, thereby significantly boosting the detection performance in future recordings. Likewise, comprehensive mechanical models of the cellular deformations due to the voltage-induced tension change provide a basis to interpret and refine the retinal recordings, including template-matching for robust detection of such weak signals.

## Conclusions

Interferometric recordings of the electrical activity in cells provide a non-invasive and label-free alternative for physiological characterization at single-cell resolution in-vivo. Optoretinography has the potential to replace electroretinography, and also greatly improve its spatial resolution. In particular, pOCT may provide co-registered structural and functional characterization of the retina. Our model of the fast contraction of the outer segment after light stimulus matches the experimental observations. The fast response is driven by the ERP, which is coupled to mechanical movement via the voltage-dependent membrane tension, leading to lateral expansion and axial contraction of the disks until the water influx takes over. The disk membrane’s very soft, undulating initial state allows significant expansion, until the membrane begins to flatten and becomes more resistant to further deformation. This leads to a logarithmic saturation of the amplitude and earlier arrival of the peak deformation with increasingly bright stimuli. Combination of this model with an accurate description of the slower phases originating in the LRP and osmotic swelling can be used in the future to characterize various aspects of the photoreceptor physiology in health and disease.

## Acknowledgements

Funding was provided by NIH grants U01 EY025501, R21EY027941, R01EY029710, P30EY001730, an unrestricted grant from the Research to Prevent Blindness, a Research to Prevent Blindness Career Development Award, the Burroughs Welcome Fund Careers at the Scientific Interfaces, and the Murdock Charitable Trust.

We would like to thank Dr. Denis Baylor for very helpful discussions about photoreceptors and the early receptor potential.

VPP, RS, and DP have a commercial interest in a provisional US patent describing the line-scan pOCT system.

## Supplementary Figures

**Supplementary Figure 1.**
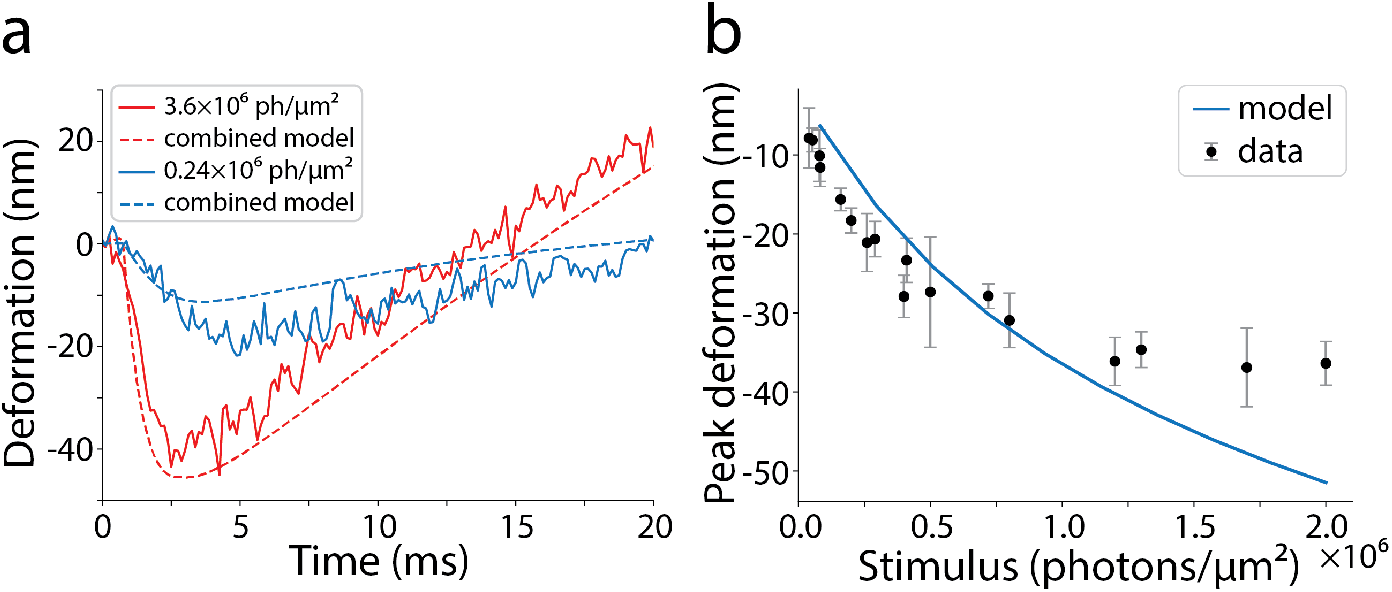
While the quasi-static model predicts roughly the correct amplitude when fit within the physiologically relevant parameter ranges, the overall fit to the measurements is poor. (a) Timing of the peaks is fixed in the quasi-static model, which is incompatible with the measurements that show later peak at weaker stimulus. (b) Saturation of the amplitude with stronger stimuli is not fast enough to fit the experimental data.

**Supplementary Figure 2.**
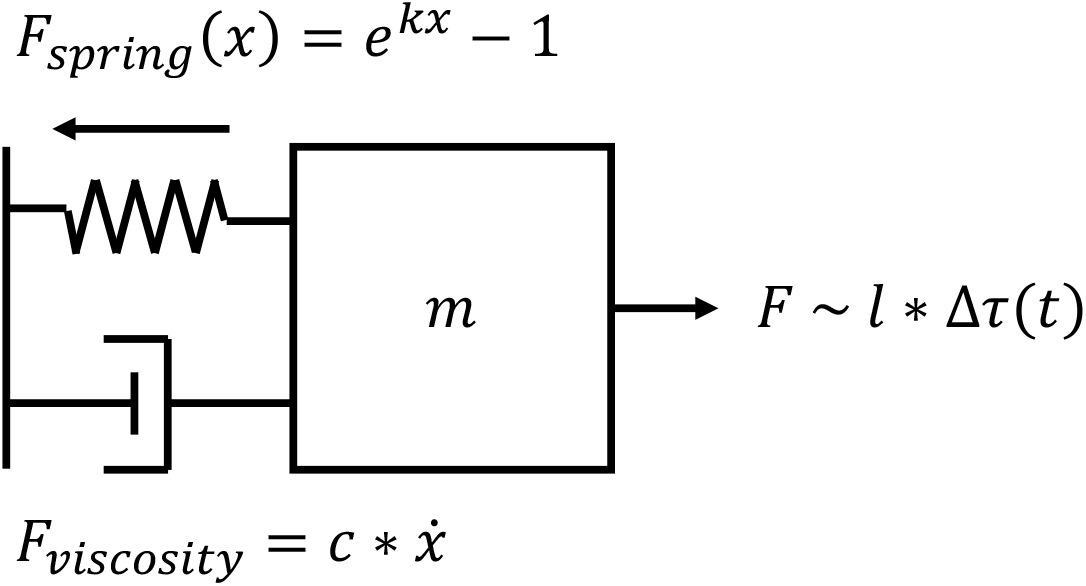
The logarithmic expansion of the membrane with increasing tension suggests a model of a mass driven in a viscous medium by a spring with a restoring force proportional to its tension, i.e. exponentially related to its deformation. The driving force represents the voltage-dependent membrane tension induced by the light stimulus.

